# CIRPIN: Learning Circular Permutation-Invariant Representations to Uncover Putative Protein Homologs

**DOI:** 10.1101/2025.11.18.689110

**Authors:** Aiden R. Kolodziej, S. Mazdak Abulnaga, Sergey Ovchinnikov

## Abstract

Protein structure-based homology detection has been revolutionized by deep learning methods that can rapidly search massive databases. However, current structural search tools often miss proteins related by topological rearrangements, particularly circular permutation (CP), where proteins share identical global folds but differ in the positioning of their termini. We introduce a circular permutation-invariant graph neural network (CIRPIN) that addresses this limitation through a novel data augmentation strategy using synthetic circular permutations (synCPs). We demonstrate that CIRPIN learns representations of proteins that are invariant to circular permutation, enabling it to identify similar proteins within the Structural Classification of Proteins - extended (SCOPe) and AlphaFold Cluster Representatives (AFDB-ClustR) databases. Leveraging the speed of CIRPIN and the accuracy of traditional structural alignment tools, we search these databases and uncover thousands of novel protein pairs related by circular permutation. Notably, we discover that PDZ domains exist naturally in four circularly permuted forms. These results highlight CIRPIN as a powerful tool to investigate the emergence of circular permutations in nature.

## 1 Introduction

Protein structure prediction tools such as AlphaFold [16] and ESMFold [24] have revolutionized biology by providing abundant structural data for drug design, function prediction, and evolutionary analysis [4]. However, these applications depend on tools to search protein structure space. While recent methods, such as Foldseek [33] and TM-vec [10] achieve remarkable speed across massive databases like the AlphaFold database (AFDB), they are limited in their ability to recover proteins related by circular permutation.

Circular permutation (CP) describes a structural relationship between two proteins wherein the global structures are the same, but the N and C termini are located at different positions. For a given protein, linkage of the N and C termini, followed by cutting the circularized protein at a new position results in a circular permutation. This rearrangement alters the protein’s contact order, making CP a valuable tool for probing protein folding mechanisms and determinants of protein stability [11, 15]. Consideration of circular permutation has also revealed evolutionarily related proteins that may have Preprint. otherwise escaped detection through conventional sequence analysis. Notable examples include the evolutionary connection between the bacterial ribosomal protein bL33 and its archaeal/eukaryotic homolog eL42 [19], as well as the relationship discovered between two carbohydrate-binding proteins [26]. These discoveries highlight the need for sensitive methods to detect circular permutations and other complex structural relationships.

While several methods exist for detecting circularly permuted proteins (reviewed in [2]), many suffer from computational inefficiency for large-scale database searches, and report scores that make distinguishing true positives from false positives difficult [7, 25, 23]. Given these drawbacks, we sought to develop a method to rapidly discover proteins related by circular permutation with high confidence.

One recent structure search method, Progres (Protein Graph Embedding Search) [9], leverages a graph neural network [29] architecture to learn an embedding space for protein structure that can be subsequently used for search. After training, a query protein can be passed through the model and searched against a pre-embedded database (Fig. 1B).

**Figure 1.**
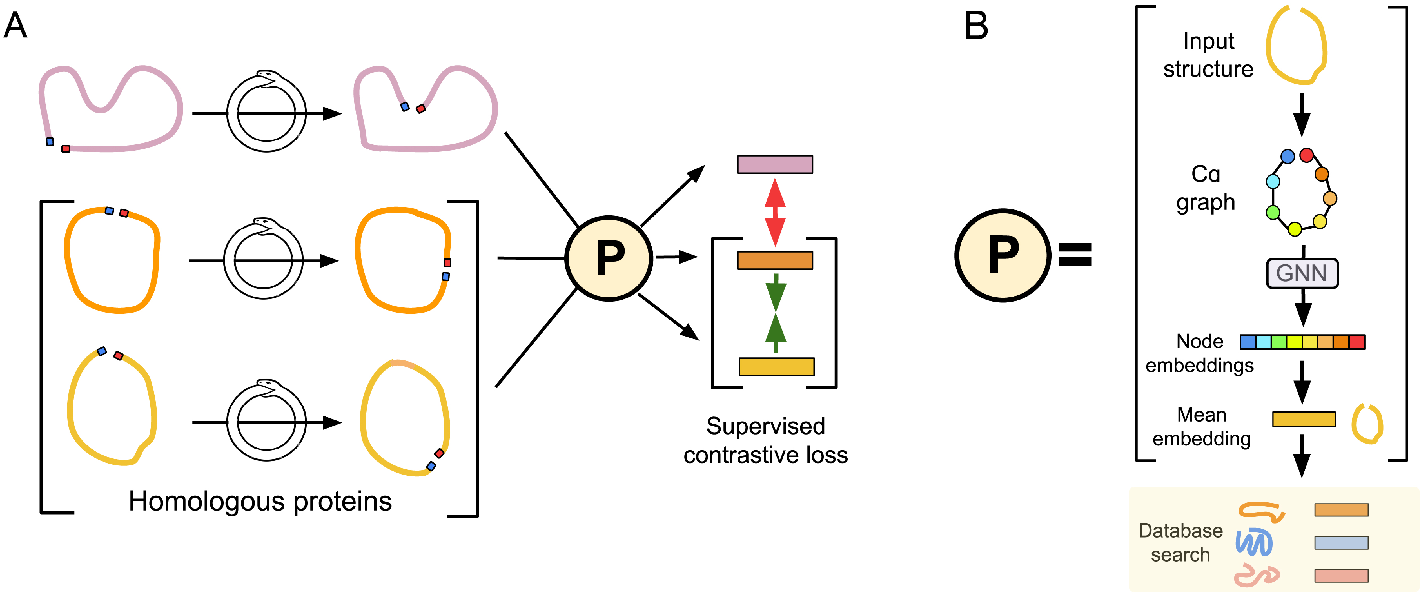
Schematic of CIRPIN and Progres model architecture. (A) `synCPs` of input structures, with N and C termini highlighted in blue and red, respectively, are generated by CIRPIN each iteration. Structures are passed through the Progres GNN architecture and a supervised contrastive loss is applied to bring embeddings of proteins in the same SCOPe family (orange and yellow) together while pushing apart proteins from other families (purple). (B) Progres model architecture. Protein structure is represented as a graph, by extracting C*α* coordinates, which is then passed through an equivariant graph neural network to obtain embeddings. Mean embeddings are then used to search for similar proteins by computing the cosine similarity between the query protein and entries in a pre-computed database. Color corresponds to positional information of each node (blue: N terminus, red: C terminus).

Building on the work of the Progres model, we introduce a **cir**cular **p**ermutation **i**nvariant graph neural **n**etwork, (CIRPIN), that results in circular permutation-invariant protein representations. By leveraging both models—one sensitive to circular permutation (Progres) and one invariant to it (CIRPIN)—we can identify structures related by circular permutation that were previously obscured by different SCOPe classifications. Notably, this dual-model approach uncovers two previously unreported PDZ domain topologies arising from circular permutation. Leveraging Progres/CIRPIN’s speed, we perform a previously intractable pairwise search over the AlphaFold Cluster Representatives Database (AFDB-ClustR) [1] and uncover thousands of previously undetected proteins related by circular permutation.

## 2 Methods

To detect proteins related by CP, we introduce a learning framework built on an existing graph neural network architecture with our proposed synthetic circular permutation operation (synCP). Our method, CIRPIN, learns embeddings that are robust to permutations in sequence connectivity while preserving essential 3D structural information of proteins. In this section, we describe how proteins are represented, formalize our synCP operation, and describe the training, evaluation, and search pipelines.

### 2.1 Protein Structure Representation

We represent each protein structure as a graph *G* = (*V, E*) where the vertices correspond to C__*α*__ atoms extracted from the protein backbone and the edges are constructed based on spatial proximity between C__*α*__ atoms. Specifically, we connect two nodes *v*__*i*__ and *v*__*j*__ with an undirected edge (*v*__*i*__, *v*__*j*__) ∈ *E* if their corresponding C__*α*__ atoms are within 10Å of each other.

For a protein containing *n* residues, we define the ordered sequence of C__*α*__ coordinates as 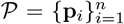, where **p**__*i*__ ∈ ℝ^3^ denotes the spatial coordinates of the *i*-th C__*α*__ atom. The indexing follows the natural N-to-C terminal directionality of the protein chain, such that **p**__1__ corresponds to the N-terminus and **p**__*n*__ to the C-terminus.

As described in [9], each vertex *v*__*i*__ ∈ *V* is associated with a feature vector **f**__*i*__ = [*ρ*__*i*__, *τ*__*i*__, **s**__*i*__] that encodes both local structural properties and sequential information where:

- 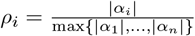 for *α*__***i***__ = { **p**__***k***__ : ∥ **p**__***k***__ − **p**__***i***__ ∥__2__ ≤ 10Å} is the normalized local density within a 10Å neighborhood.
- 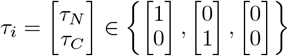 where the vectors correspond to whether it is N-terminal, C-terminal or neither.
- **s**__*i*__ ∈ R^64^ is a sinusoidal positional encoding [34].

### 2.2 Synthetic Circular Permutation

To test a model’s performance on identifying similarity between proteins related by circular permutation, we introduce the concept of synthetic circular permutations (synCPs). For any protein of length *n*, there are *n* possible circular permutations. We generate synthetic circular permutations of a given protein by circularly reordering the protein’s C__*α*__ coordinates.

Given the original ordered sequence of C__*α*__ coordinates 𝒫, a *k*-synthetic circular permutation is defined as:

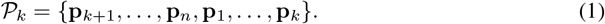

This operation can be implemented in PyTorch using *𝒫*__*k*__ = torch.roll(𝒫, *n* − *k*).

Two structures related by synCP will have nearly identical graphs, with edges differing only at the original and new N and C-termini due to the breaking and formation of terminal connections. However, the positional embeddings **s**__*i*__ at each node will be circularly permuted according to the new sequence ordering. The terminal indicator feature *τ*__*i*__ will also change to reflect the new termini positions. Due to the fact that these graphs do not necessarily correspond to real circular permutations that occur in nature, we refer to them as “synCPs.”

### 2.3 CIRPIN Architecture and Training

CIRPIN builds upon the Progres model architecture, which uses an E(n)-equivariant graph neural network [29] and supervised contrastive loss to embed protein structures [17] (see Fig. 1B).

We add a synCP generation module that creates random circular permutations during training (Fig. 1A). We extend the construction of positive and negative pairings of proteins described in [9], where positive pairings are computed by taking proteins in the same SCOPe family. In each training iteration, we sample a structure and perform a random synCP on that structure. This operation is represented by the ouroboros, an ancient symbol of rebirth, on Fig. 1A. Over the course of many epochs, the model sees a diverse range of synCPs of each protein in the training set. Using supervised contrastive loss, proteins within the same SCOPe family are brought closer together in latent space and more distantly related proteins are moved far apart. However, upon implementing CIRPIN, distantly related proteins sharing higher SCOPe classification, such as fold or superfamily, may now be pushed closer in latent space.

We trained the model for 500 epochs and used the Progres validation set to monitor performance on recovering domains in SCOPe from the same fold, superfamily and family, as described in [33]. We selected the model with best validation set performance for all further use. After training the model, database search is performed rapidly by ranking the cosine similarity between the embedding of an input protein against all proteins in a pre-embedded database.

### 2.4 Structural Alignment with TM-align

We use TM-align, a structural alignment algorithm based on the TM score metric [36] and commonly used as a benchmark metric in competitions such as CASP [20] as our reference structural alignment method. A TM score of < 0.17 is considered to represent random or no significant structural similarity, whereas a score > 0.5 suggests the proteins are highly similar as it would take at least 1.8 million random structural matches so that one structure match reaches a TM-score ≥ 0.5 [36]. We use ≥ 0.5 as a cutoff for CP retrieval. TM-align returns two scores, normalized by either the query or target length. In all analyses we take the minimum score returned by TM-align, since we seek a score reflecting global similarity between two structures.

TM-align depends upon dynamic programming to generate optimal alignments. Consequently, the algorithm is sensitive to sequence order, and thus to aligning circular permutants. However, an optional -cp duplicates the query structure to allow the alignment to wrap around, enabling detection of circular permutations. Variations of this general approach, “duplicate and align”, have been used in several previous works [32, 3, 25, 13].

Since TM-align CP also reports high scores for protein pairs *unrelate*d by CP, it is necessary to compute both the TM-align and TM-align CP score. By taking the difference, Δ*CP*, we can identify pairs in which consideration of circular permutation resulted in an improved alignment. We calculated the Δ*CP* score for the test set of CPs and found that pairs of CPs had the highest Δ*CP* scores (Fig. S2).

## 3 Results

We first assess the ability of Progres and TM-align [36] to find proteins related by CP. Using our defined synCP operation we probe the dependence of these two methods on positional information and find they have a limited ability to detect known CPs. We then compare our model, CIRPIN, against Progres on retrieving known example CPs from the literature. Subsequently, we use CIRPIN to identify novel CPs in the SCOPe and the AFDB-ClustR. Finally, we demonstrate the learned embeddings of CIRPIN capture structurally meaningful features and CIRPIN can be additionally used to recover similar proteins obscured by secondary structure rewiring and insertions.

### 3.1 Probing Structural Alignment Methods Using SynCPs

Given that embeddings across various lengths are averaged in Progres, instead of aligned, we postulated Progres embeddings of protein structure might be invariant to sequential perturbation, such as circular permutation, and could be used to identify proteins related by CP. To investigate this idea, we evaluated whether Progres can detect proteins related by CP within the SCOPe database (clustered at 40% sequence identity) [8].

As a representative example, we focused our attention on the winged helix-turn-helix (wHTH) fold, a widely conserved DNA binding domain found in transcription factors across the tree of life [21]. Previous structural characterization of a polysaccharide biosynthesis protein from *C. acetobutylicum* revealed the presence of a circularly permuted wHTH [6]. These two structures are also present in the SCOPe database as the rpc34 subunit in RNA polymerase III from mouse, d2dk8a1 (hereafter referred to as “K8”), and as the human methionine aminopeptidase d1qzya1 (hereafter referred to as “QZ”). Embedding these two domains using Progres and evaluating their homology via cosine similarity returned a low score of 0.39 (Fig. 2A), which following a similarity threshold of 0.8 [9], indicates no significant structural similarity.

**Figure 2.**
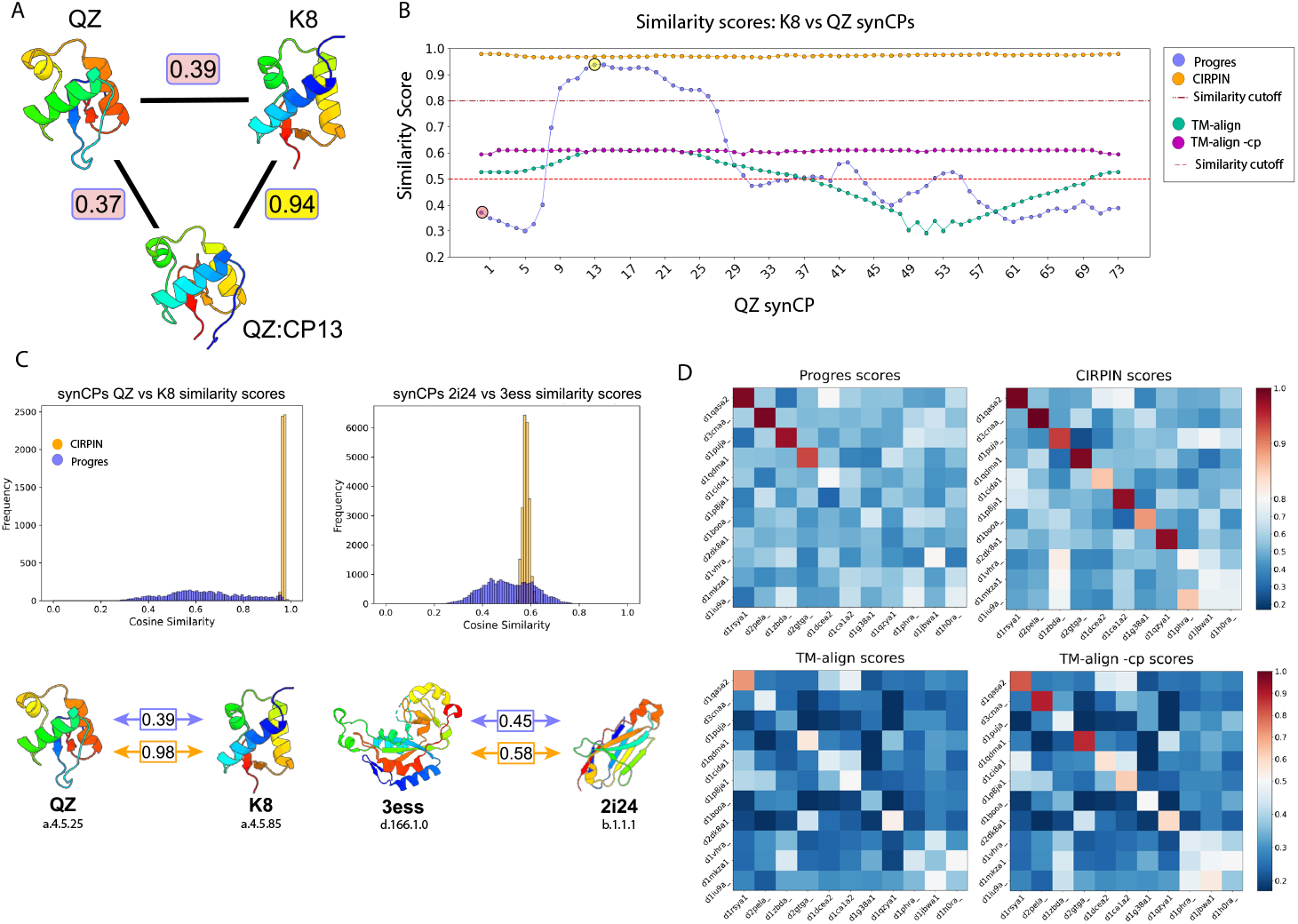
Evaluation of structure search methods on retrieval of proteins related by circular permutation. (A) Scores between QZ, K8, and the 13th synCP of QZ. Progres scores highlighted in yellow (high) or red (low). Structures colored from N (blue) to C terminus (red). (B) Similarity scores between every synCP of QZ vs unpermuted K8. Horizontal lines mark the similarity cutoff score for Progres/CIRPIN (0.8) and TM-align (0.5) (C) All versus all comparison of query synCPs vs target synCPs. Scores between unpermuted original structures from the two models shown below. SCOP(e) concise classification string (sccs) shown below structures, indicating *class*.*fold*.*superfamily*.*family*. (D) Benchmark set of verified pairs of circular permutations arranged such that scores for pairs of CPs follow the diagonal of the heatmap. Heatmap colors are centered at the similarity cutoffs of 0.5 and 0.8.

We hypothesized that creating a synCP of a query structure such that the node ordering aligns to the amino acid order of a circularly permuted target protein would result in detection by Progres without any retraining of the model. To test this idea, we generated all possible synCPs for query QZ and scored them against the target K8. The results show a wide range in scores that is solely dependent on amino acid ordering (Fig. 2B). As predicted, some synCPs of QZ now score highly against K8 (Fig. 2B). The highest scoring structure was QZ permuted at position 13 and aligns well to K8 (Fig. 2B).

We also investigated if TM-align could identify similarity between these two proteins. As shown in Fig. 2B, although TM-align found similarity between the unpermuted two structures, it struggled to do so for many synCPs (33 out of 73). However, with the -cp flag implemented, the resulting scores are unaffected by circular permutation and are all above the 0.5 fold similarity cutoff (Fig. 2B). However, it is well established that TM-align and related tools are far too slow to use on the size of today’s structural databases. For example, the authors of Foldseek [33] estimated that using TM-align to calculate an all vs all comparison of 100 million structures would take approximately 10 millennia on a 1,000 core cluster. This issue is further exacerbated by the fact that running TM-align -CP is ∼ 55% slower than running TM-align.

Next, we generated all synCPs for both the query and target and scored them in an all-versus-all manner, resulting in 5, 920 comparisons for QZ and K8. Plotting the Progres scores of all possible synCP comparisons produced a wide distribution of scores (Fig. 2C), indicating that the score between two proteins can vary greatly, depending solely upon the amino acid order. We found this remained true for two random proteins unrelated by CP and which have different SCOPe class designations (Fig. 2C). We also investigated whether TM-vec [10] has a similar limitation and found that scores between synCPs can vary considerably (Supplemental Fig. S1).

Based on these results and motivated by Progres’s speed relative to TM-align, we investigated how Progres could be modified to achieve circular invariant protein representations. To this end, we developed CIRPIN.

### 3.2 Benchmarking CIRPIN

We hypothesized that by training the Progres graph neural network (Fig. 1b) on our proposed synthetic circular permutants (synCPs), the model would learn embeddings such that circular permutants are scored as similar. Furthermore, such a model would become less dependent on positional information and instead learn interacting structural motifs to more accurately embed proteins in a shared latent space. We first considered removing positional information entirely, but as shown in [9], this drastically reduces the model’s ability to accurately retrieve homologous proteins.

After training CIRPIN, we used our model to embed QZ and K8 and found CIRPIN correctly scored them as highly similar (Fig 2C). We then compared the distribution of CIRPIN and Progres scores (Fig. 2C) for all synCPs of QZ and K8. The narrow distribution of CIRPIN scores indicates that circularly permuting the positional information, by generating synCPs, has little effect on the score calculated by CIRPIN, whereas it greatly alters the scores output by Progres. These observations held true when comparing two unrelated proteins (Fig. 2C).

#### 3.2.1 Retrieval of Homologous Proteins

One concern with training on synCPs is that CIRPIN might overgeneralize and identify similarity in structurally dissimilar proteins. As a sanity check, we evaluated CIRPIN’s ability to recover domains from the same SCOPe fold, superfamily, and family using the test set constructed by Progres. This set comprises 400 randomly selected domains from the Astral 2.08 40% sequence identity dataset, with no training set domains sharing ≥ 30% sequence identity with these test cases. Table 1 shows that CIRPIN achieves recovery scores comparable to Progres (higher is better). We expect CIRPIN to perform slightly worse than Progres on this task, since CIRPIN is trained, by means of our synCP operation, to learn similarity between proteins that are not already neatly classified into the same family by SCOPe.

**Table 1:** Comparison of ability to retrieve homologous proteins from SCOPe.

| Model | Fold | Superfamily | Family | SCOPe Database Run Time (all vs all) |
| --- | --- | --- | --- | --- |
| CIRPIN | 0.166 | 0.692 | 0.865 | 154 seconds |
| Progres | 0.177 | 0.706 | 0.877 | 155 seconds |

#### 3.2.2 Recovering Circular Permutations in SCOPe

An ideal measure of CIRPIN’s performance at recovering protein pairs related by CP would be testing using a benchmark dataset consisting of structurally diverse pairs of circular permutants. Unfortunately, no such set has been established in the field, with researchers relying on a only a few well-studied domains, such as the concavallin A lectin domain [27, 28, 13] to test their methods. To address this issue, we performed a thorough literature review of reported circular permutants and constructed a test set of 11 protein pairs which were aligned in PyMOL [30] and verified visually to be related by CP.

We scored this database using TM-align (see Methods): TM-align recovered 3 out of 11 and TM-align -cp recovered 7 out of 11 (Fig. 2D). For retrieval using Progres or CIRPIN, a cosine similarity above 0.8 corresponds to a successful recovery of a homolog [9] (Fig. 2D). Of the 11 test domains, Progres recovered 4 and CIRPIN recovered 9. However, two of these pairs are grouped within the same SCOPe family, and so during training the embeddings of these structures would have been pushed closer to one another. Since we seek to determine how well the two models perform at recovery in instances where SCOPe does not classify circular permutants in the same family, the adjusted recovery rates are 2 out of 9 for Progres and 7 out of 9 for CIRPIN (Fig. 2D). The two pairs CIRPIN was unable to identify were also missed by TM-align -cp, indicating that other structural changes, in addition to circular permutation, obscure the structural similarity between these proteins. For example, a careful visual comparison of d1iu9a and d1h0ra revealed that a region composed of a long beta sheet in d1h0ra is present as a shorter beta sheet and alpha helix in d1iu9a.

### 3.3 Identifying Novel Circular Permutations in SCOPe

We embedded the entire SCOPe database (clustered at 40% sequence identity, 15.2k structures) using both CIRPIN and Progres models, then performed an all-vs-all comparison. To identify pairs related by circular permutation with high confidence, we developed the search pipeline outlined in Fig. 3A. We use Progres and CIRPIN as a rapid filter for identifying putative CPs and then use TM-align as a sensitive alignment method to verify these hits.

**Figure 3.**
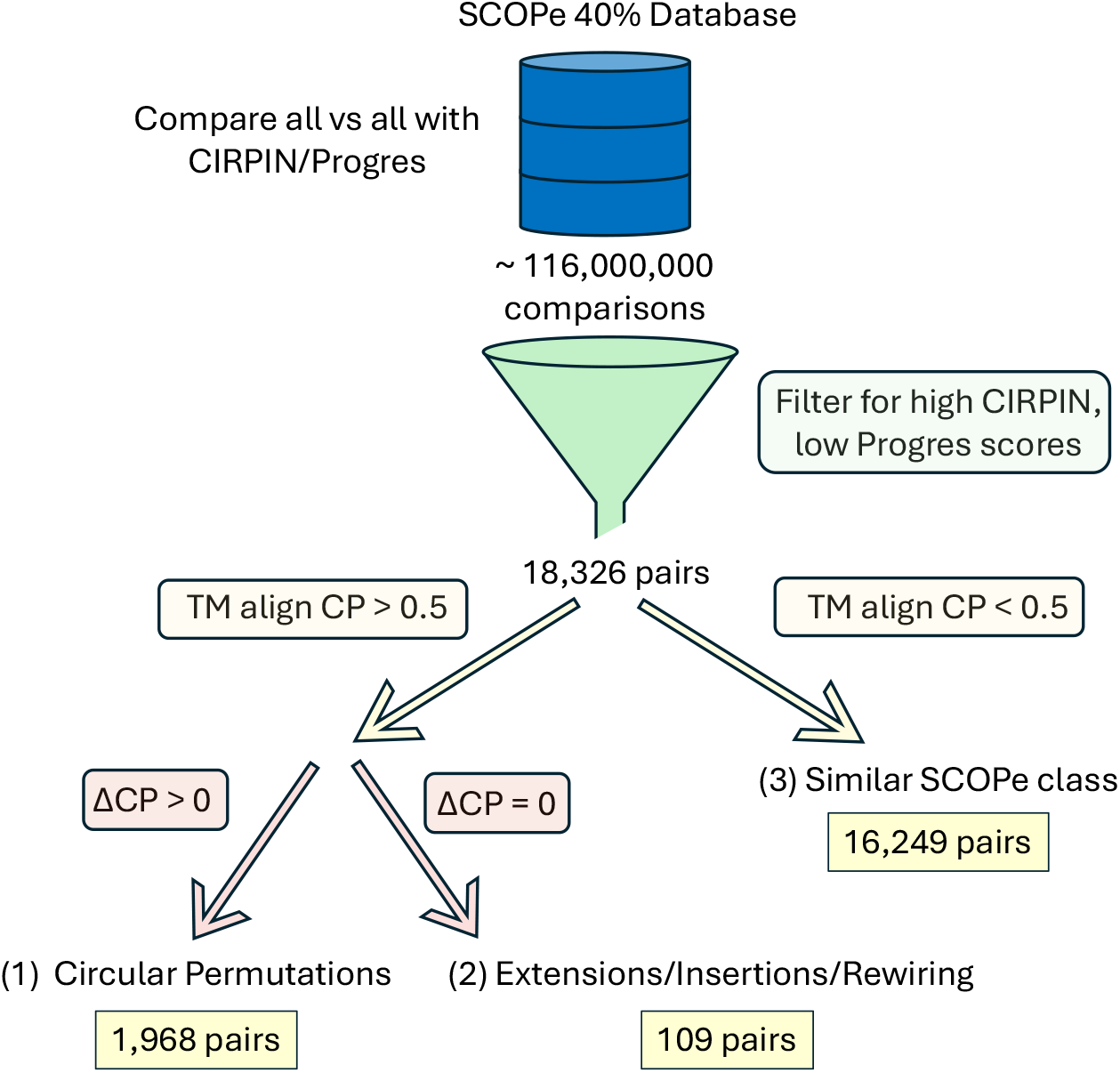
Schematic for circular permutation search. Cutoffs used for Progres and CIRPIN scores were 0.6 and 0.9, respectively. ΔCP is the difference between TM-align and TM-align -cp scores.

Following filtering, we obtained 18, 326 protein pairs with high Progres/CIRPIN score differences. Most of these 16, 249 pairs (∼ 89%) had TM-scores below 0.5, suggesting different folds and likely excluding cases of circular permutation. Further analysis revealed that pairs in this bin tend to have similar secondary structure content. Specifically, pairs in this bin share the same SCOPe class ∼ 80% of the time, a 4-fold enrichment over the ∼ 20% baseline chance of two random proteins sharing the same SCOPe class.

Using the difference between TM-align with and without a CP flag (see Methods) we identified 1, 973 pairs related by circular permutation. Accounting for duplicates, there are 1, 204 unique structures. This suggests that approximately 8% of structures within the SCOPe database are related to another structure by CP.

To investigate the structural diversity of the identified CPs, we grouped them by the same SCOPe class and fold and plotted the number of occurrences (Fig. 4A, Fig. S3). We then examined each class and identified folds that were overrepresented. An analysis of the beta class showed folds previously known to contain circular permutations (b.7, b.12, b.18) and revealed b.36 (PDZ domain-like fold) as a fold containing a high number of CPs (Fig. 4A). Based on these results, we further investigated the structures within the PDZ domain-like fold.

**Figure 4.**
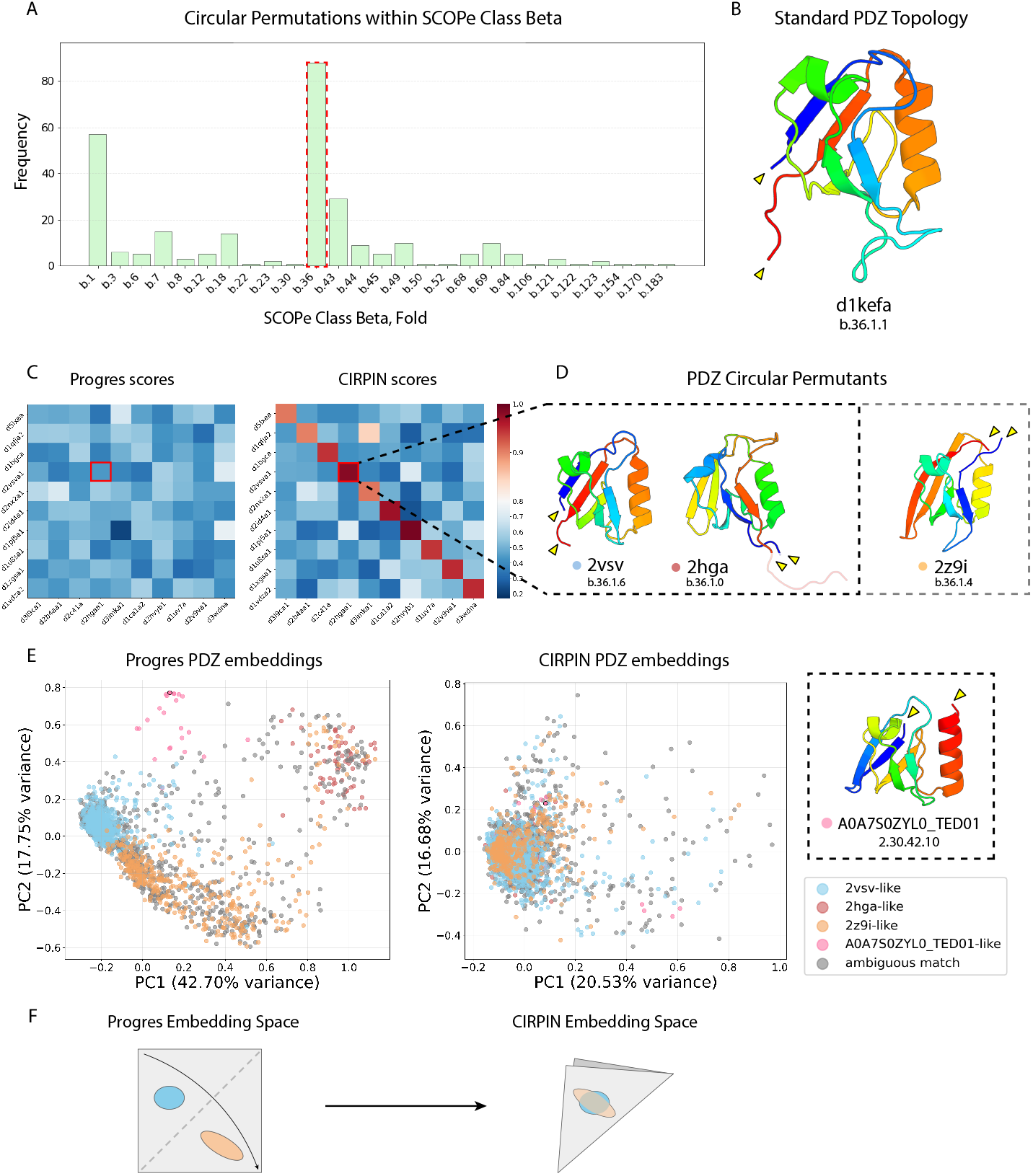
Discovery of novel circular permutations of PDZ domains by CIRPIN. (A) Schematic for circular permutation search. Cutoffs used for Progres and CIRPIN scores were 0.6 and 0.9, respectively. ΔCP is the difference between TM-align and TM-align CP scores. (B) Number of circular permutations found in folds within the SCOPe class beta. Red highlights PDZ domain-like fold, b.36. (C) Heatmap of Progres/CIRPIN scores between representative protein pairs discovered through (a). (D) Structures of three PDZ domains related by circular permutation, colored from N to C terminus. (E) PCA of AFDB-ClustR PDZ domains. Novel topology discovered in AFDB-ClustR colored in pink, CATH label displayed below. (F) CIRPIN “folds” the Progres embedding space to bring together structures related by CP.

#### 3.3.1 Novel Circular Permutations of PDZ Domains

PDZ is an acronym for three proteins first discovered to share the same domain: post synaptic density protein (PSD95), Drosophila disc large tumor suppressor (Dlg1), and zonula occludens-1 protein (zo-1). We examined PDZ domains from these three proteins and found they all shared the same N and C termini locations. We show one of these structures, the PDZ domain from PSD95/SAP90 (d1kefa) as the “standard” PDZ topology since most research has focused on PDZ domains exhibiting this topology (Fig. 4B).

Among the CPs identified by our search within the PDZ domain-like fold are two PDZ domains related by CP, PDB: 2vsv and PDB: 2hga. While previous research identified a circular permutation of a PDZ domain in *S. obliquus* [14] represented by the 2z9i structure, CIRPIN identifies an additional permutant, represented by the 2hga structure (Fig. 4D). We noted that 2hga contains a long C-terminal tail, which led us to wonder whether this feature contributed to its obscuration from Progres. However, truncating the C-terminal tail of 2hga did not improve Progres’s ability to detect homology between 2hga and 2z9i or 2vsv, while CIRPIN continued to assign a high score. For clarity in comparing the three topologies, we present the truncated version of 2hga. Comparison of the domain found in this study and [14] reveals that there exists at least three distinct topological variants of PDZ domains, each related to one another by CP (Fig. 4D). These three domains are classified into separate SCOPe families, partly explaining Progres’s failure to retrieve them: the model’s training objective explicitly optimizes for high intra-family similarity and low inter-family similarity.

Given our discovery of a novel PDZ topology within SCOPe we asked whether additional topologies may exist in larger databases. To address this, we took The Encyclopedia of Domains (TED) [22] annotated PDZ domains from the AFDB-ClustR (4, 015 domains) and filtered for PDZ domains between 75-100 amino acids in length in order to remove poor quality annotations. This yielded 2, 563 structures. We then plotted a PCA of the Progres and CIRPIN embeddings of these structures (Fig. 4E).

While analyzing the Progres PCA plot, we noticed a cluster of structures that appeared distinct from the three reference structures (Fig. 4D) we found in SCOPe. A representative structure from this cluster, shown in (Fig. 4E) constitutes another distinct PDZ circular permutant. We then colored the PCA plot according to which reference PDZ structure they best aligned to using TM-align. In some cases, a domain aligned well to multiple reference structures. If these scores were within 0.025 of each other, we assigned these domains an “ambiguous match” label.

Owing to its circular permutation invariance, CIRPIN embeds CP-related proteins close to one another in space. Relative to Progres, this can be viewed abstractly as “folding” the Progres embedding space, whereby proteins related by circular permutation, once separated in space, now cluster together (Fig. 4F). By comparing the embeddings of a circularly invariant model (CIRPIN) to a non-circularly invariant model (Progres) we were able to both discover new circular permutations and investigate the number of distinct circularly permuted topologies.

### 3.4 Identifying Novel Circular Permutations in AFDB-ClustR

After verifying that our method can recover novel CPs from SCOPe, we applied the method to search for CPs in the AFDB-ClustR [1]. We first processed the AFDB-ClustR into individual domains using Foldcomp [18] and The Encyclopedia of Domains (TED) annotations [22]. Since this database is over two orders of magnitude larger than SCOPe, we used a more stringent filtering citeria; we filtered for pairs that have a Progres/CIRPIN score difference of 0.7. This resulted in 833, 387 pairs, which we subsequently verified using TM align/TM align -cp following the schematic shown in (Fig. 4A).

We then filtered for pairs that had a TM-align -cp score > 0.5 and a Δ CP score > 0. From this dataset we identified 137, 020 unique domains that have at least one structure related by circular permutation within the AFDB-ClustR. Among the most represented domains are those without CATH annotations, Winged helix-like DNA-binding domains (1.10.10.10) and Ubiquitin-like (UB roll) domains (3.10.20) (Fig. 5A).

**Figure 5.**
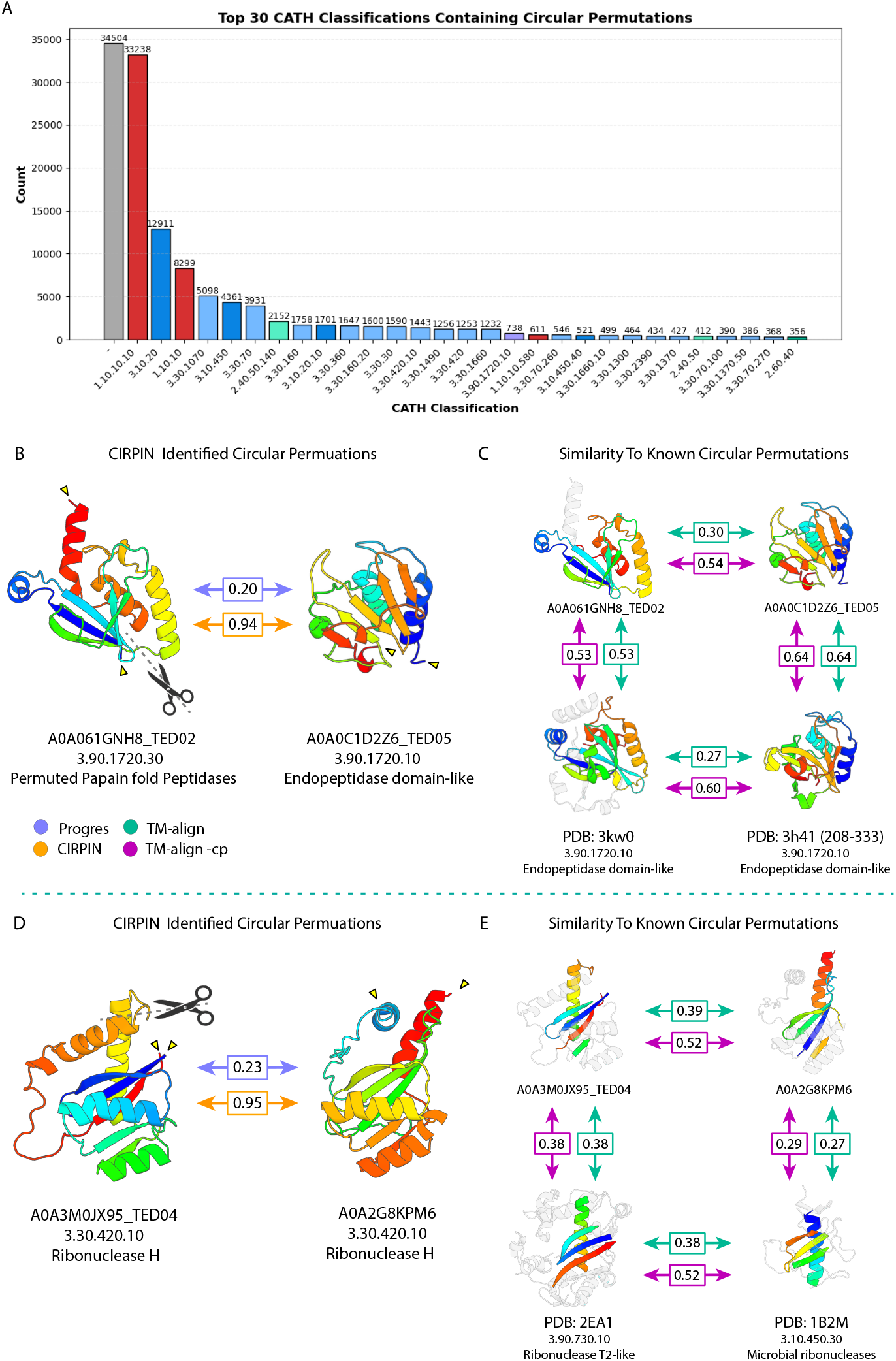
Circular permutations in the AFDB-ClustR. (A) Distribution of CP structures according to TED CATH annotations (colored by CATH architecture). (B) Endopeptidase domains related by CP. (C) Comparison of domains in (b) to a previously identified pair of endopeptidases. (D) Ribonuclease H-like domains related by CP.(E) Comparison of domains in (d) with functionally related ribonuclease T2-like domains. Labels show the CATH level (Class, Architecture, Topology, Homologous Superfamily). Peripheral structural elements are colored gray to highlight the shared core structural motifs. Scores calculated from colored regions of the structures.

Encouragingly, we found pairs of circular permutations that corroborate previous studies. We identified a pair of circularly permuted endopeptidases with high similarity to those identified by [35] (Fig. 5B,C). Additionally, we a novel circular permutation relationship between two Ribonuclease H-like domains (Fig. 5D). Interestingly, based on its CATH annotation this pair is functionally related, though structurally divergent, to a pair of ribonuclease T2-like domains previously found to be related by circular permutation [23] (Fig. 5E).

In addition, our dataset reveals thousands of novel proteins pairs related by CP. Representative examples of these pairs are shown in (Fig. 6A). Importantly, many pairs are classified into different CATH classes, architectures, and topologies, or lack CATH annotations altogether. This suggests that simple searches based on CATH or SCOPe family labels alone are severely limited in their ability to recover proteins related by CP.

**Figure 6.**
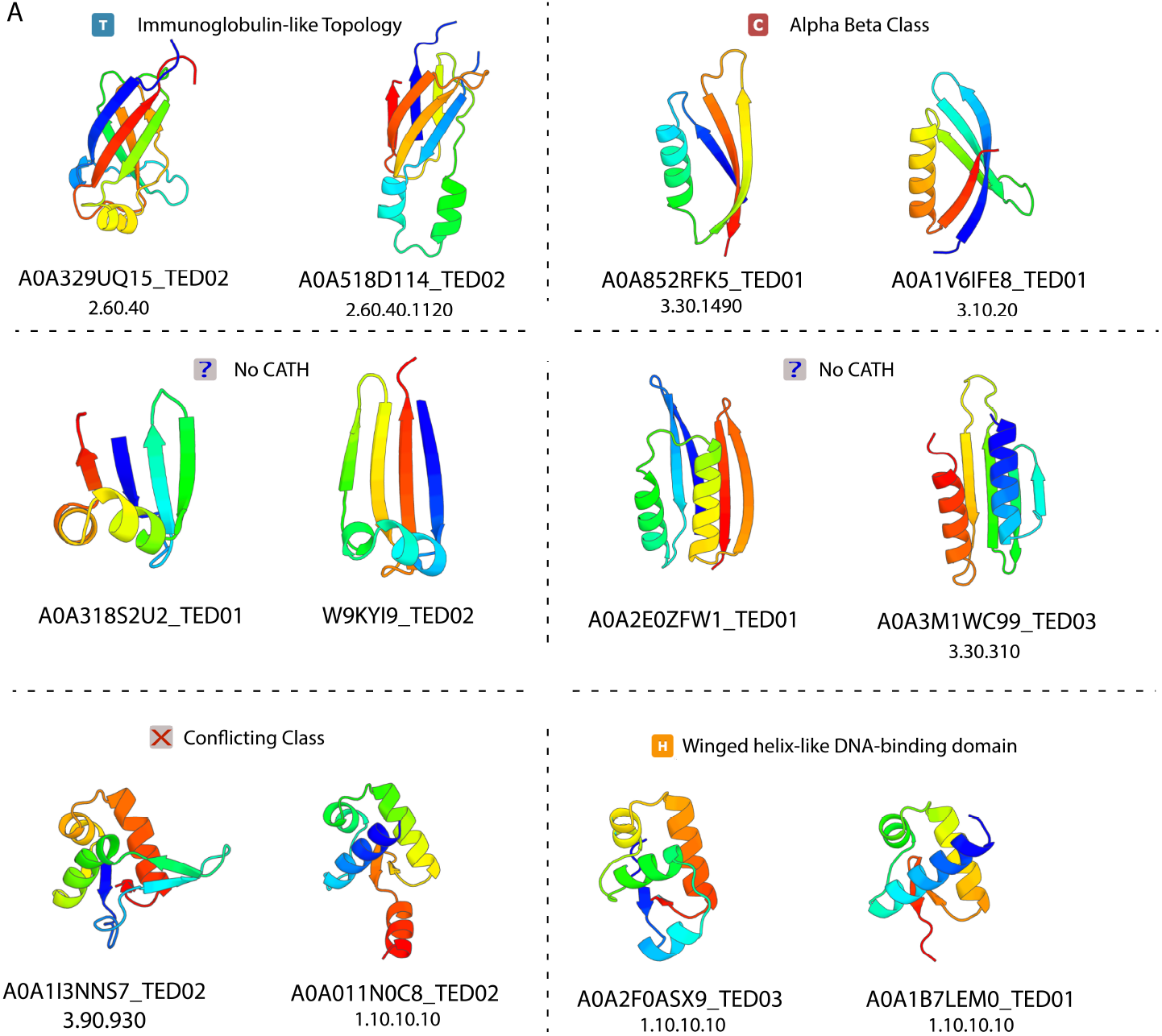
Novel circular permutations in the AFDB cluster representatives. Icons above each pair indicate the CATH hierarchy shared between the two structures.

### 3.5 CIRPIN Retrieves Protein Pairs Obscured by Rewiring and Insertions

Among the structures that had the largest difference in Progres/CIRPIN scores, we noted many that appeared not to be circular permutants of one another (ΔCP = 0) (Fig. 7), but rather contained insertions, extensions, and/or secondary structure rewiring. We hypothesized Progres embeddings may be highly sensitive to precise domain definitions. To test this, we removed the insertion from a representative example, re-scored the domains and found this significantly increased the Progres score (Fig. 7D). Furthermore, we found that these alterations resulted in negligible changes in CIRPIN scores and therefore show a single CIRPIN score for this comparison. We also noticed structures which share the same fold, but have different secondary structure connectivity (Fig. 7A,B,C). The structures of one pair, shown in (Fig. 7A), are classified in the same SCOPe family, yet Progres fails to find similarity between them. (Fig. 7B) shows a simplified diagram of these two structures to highlight the differences in connectivity. These results suggest that CIRPIN embeddings can be used to narrow the search space for pairs of proteins that may be related by more complex rearrangements.

**Figure 7.**
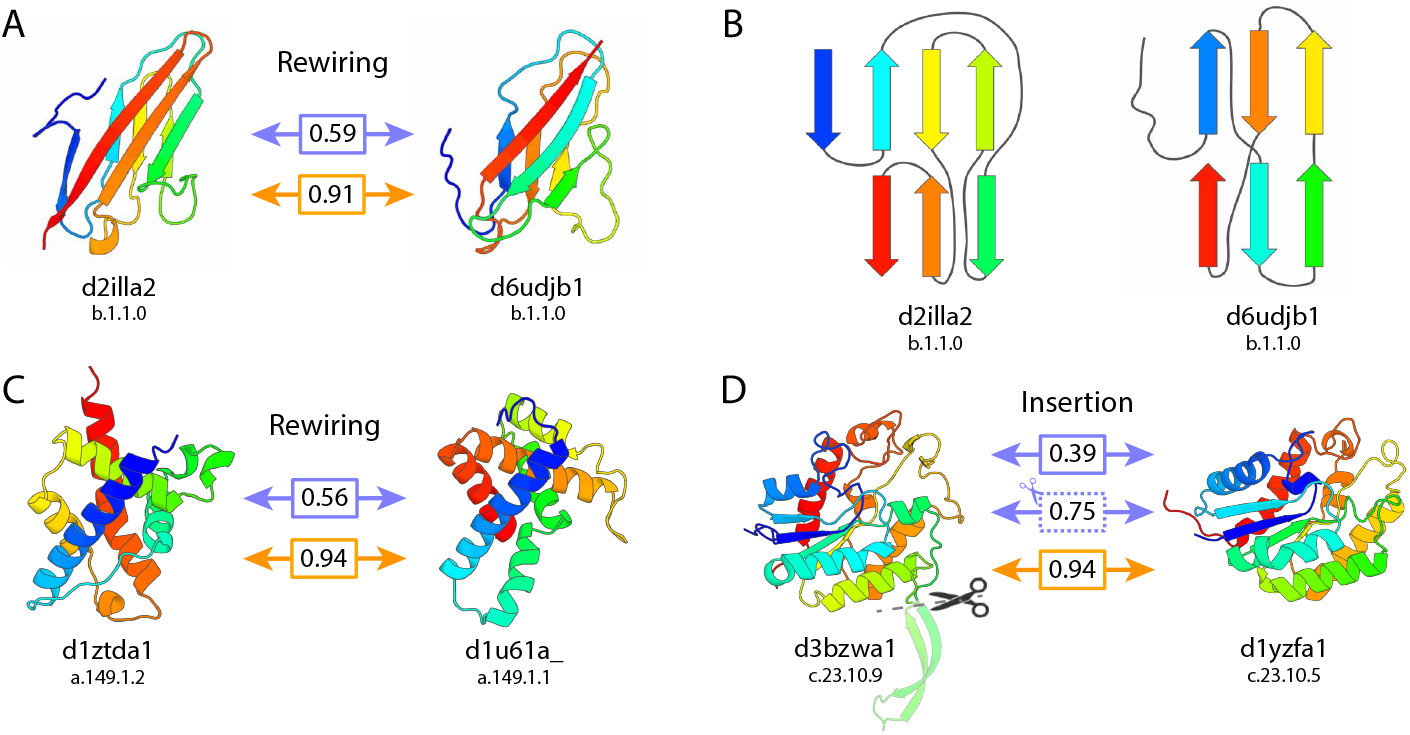
CIRPIN recovers proteins previously obscured by insertions and rewired secondary structure. (A,C) Proteins with similar architecture, different connectivity, recovered by CIRPIN. (B) Diagram showing differing connectivity of structures in (a). (D) Homologs previously obscured by an insertion. Boxes from top to bottom: (top: Progres score on default structures), (middle dotted: Progres score after removing insertion/extension), (bottom: CIRPIN score on default structures).

### 3.6 Tertiary motifs emerge from CIRPIN node embeddings

To better understand how CIRPIN learns accurate embeddings for protein structure, we investigated the per node embeddings of proteins (Fig. 8A). Here we present the analysis of the N-terminus of PDB 1uur, to more easily decipher differences between CIRPIN and Progres per node embeddings.

**Figure 8.**
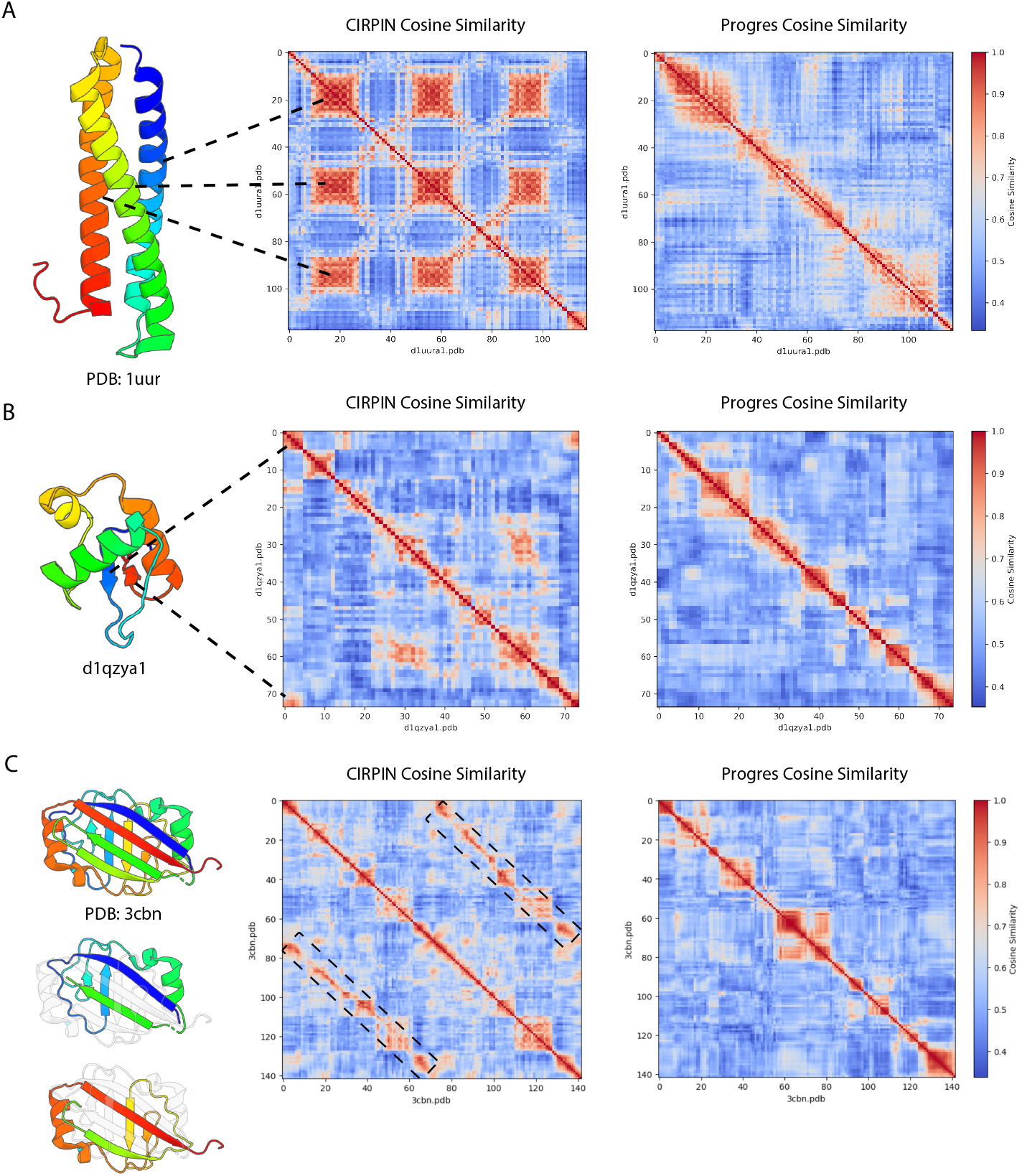
CIRPIN embeddings capture similarity between tertiary motifs that are absent in Progres embeddings.(A) CIRPIN and Progres node level embeddings for N-terminus of PDB 1uur (self vs self). Alpha helices show similarity to one another. (B) CIRPIN and Progres node level embeddings of QZ (self vs self). N and C terminal beta sheets show high similarity. (C) CIRPIN and Progres node level embeddings of PDB 3cbn (self vs self). Super-imposable substructures of 3cbn show similarity in CIRPIN node-level embeddings (Black dashed regions).

The N-terminus of 1uur consists of three alpha helices of nearly equal length connected by two loop regions. Fig. 8A shows the result of extracting the per node embeddings of 1uur and plotting the cosine similarity between all nodes. Strikingly, the node-level heatmap for CIRPIN shows three regions of high cosine similarity that closely align to the alpha helices in the structure, indicating a correspondence between protein tertiary structure and latent space representations (Fig. 8). Analysis of QZ shows that CIRPIN finds similarity between residues within the two short N and C terminal beta sheets (Fig. 8B). These patterns of structural similarity are markedly absent in the same heatmaps generated by Progres (Fig. 8B).

To show whether CIRPIN’s ability to capture similarity within domains extends beyond simple recognition of alpha helices and beta sheets, we examined the more intricate structure of PDB: 3cbn. This protein has unique internal symmetry: the structure can be parsed into two nearly identical substructures that interlock to form a single domain. These two substructures superimpose nearly perfectly (TM-score = 0.81) and are shown individually in (Fig. 8C). Analysis of the node-level embeddings show that CIRPIN identifies these two substructures as highly similar, while the Progres heatmap lacks this signal. (Fig. 8C). Taken together, these results indicate that by training on synCPs, CIRPIN learns features based on local context, while Progres may over-rely on the positional information of residues. Given CIRPIN’s ability to capture substructure similarity, these embeddings could be used in future work to obtain explicit alignments.

## 4 Discussion

We introduce CIRPIN, a deep learning model that, combined with traditional structure alignment tools, enables systematic detection of circularly permuted proteins. Using CIRPIN, we generated the largest dataset to date of protein pairs related by circular permutation: CIRPIN-DB. Our dataset greatly expands the number of known CPs and will serve as an important resource for future studies investigating the evolution of circularly permuted proteins. In addition to circular permutations, our identification of proteins related through extensions, insertions, or rewiring expands the set of remote homologs available for benchmarking challenging structure search tasks, addressing the severely limited size of current databases used for this task [5].

Among the novel pairs of CPs we identified are PDZ domains which we found to exist in four topological forms, each related by circular permutation. Although much has been published on the peptide specificity of the PDZ domain, few studies have examined the topological heterogeneity of this family. PDZ domains represent the most frequently inserted domain family [12], and characterization of their structural variants presented here will inform future work investigating the evolutionary radiation of this domain.

One important caveat of our current search method is the stringent Progres/CIRPIN score difference we used. We noticed that from our filtering criteria, no structures with the PDZ CATH label, (2.30.42.10) appeared in our AFDB-ClustR hits, indicating that our strict score difference is limiting our detection of all CPs. We anticipate by adding additional filters, or adding an explicit alignment of CIRPIN node level-embeddings during training, we can improve the model such that less drastic score differences can be used and additional pairs can be recovered. Additionally, since CIRPIN was trained on the SCOPe database, it is unlikely to generalize well to proteins that are highly divergent from those in the database. Future work can fine-tune or retrain the model on sets of novel folds.

While CIRPIN identifies proteins related by CP, it is important to note that such a relationship does not necessarily indicate that the two proteins descended from a common ancestral sequence. Two structures could converge independently to the same fold, with differing N and C termini. Future work could investigate specific examples identified by CIRPIN to unravel to the evolutionary events leading to circular permutation and to provide evidence for either convergent or divergent evolution.

Lastly, our work has important implications for how train/test splits should be constructed for any model trained on protein structure. If a model is trained on structures split by at the CATH topology level, they may unintentionally contain data leakage in the form of circularly permuted structures that appear in different CATH architectures.

More generally, our method presents a powerful use case of semi-synthetic data generation for addressing limitations of current structure search tools. While other methods for circular permutation detection have relied upon modifications to traditional alignment algorithm such as Smith-Waterman [31], our method avoids explicit alignment entirely, enabling rapid search. To encourage further investigation, we provide a complete list of CPs identified by CIRPIN and the model weights and training script at: https://github.com/aidenkoloj/CIRPIN.

## Acknowledgments

We thank members of the Ovchinnikov lab for helpful discussions, Michael Feigen for assistance on formalizing our synCP operation, Joseph L Watson for PyMOL aesthetics settings used in the figures, Joe Greener for Progres training code and training set. S.O. acknowledges support from NSF grant MCB2032259, CocaCola and Amgen. S.M.A. acknowledges support for this research by the NIH National Institute of Mental Health UM1MH130981, R01 MH123195, R01 MH121885, 1RF1MH123195 grants.

## A Supplementary Material

**Supplementary Figure S1:**
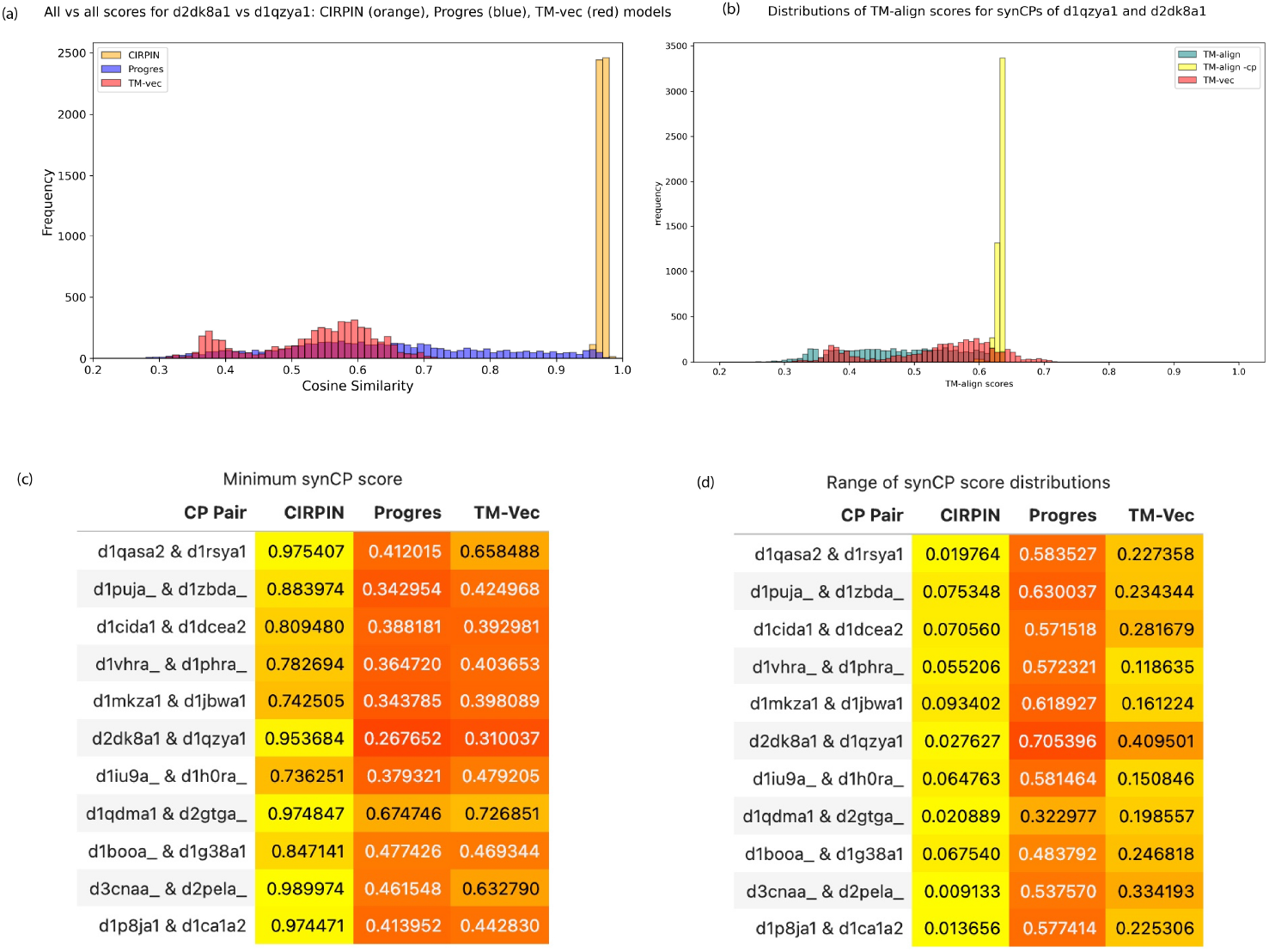
(a-d) EDITComparison of Progres [9], CIRPIN, and TM-vec [10] performance on scoring a benchmark dataset of circular permutant pairs. (a) Representative distribution of synCP scores for circular permutants QZ and K8. (b) Minimum synCP scores taken from distributions shown in (a) for all pairs of circular permutants; higher scores (indicating structural similarity) are highlighted in yellow. (c) Range of synCP score distributions for Progres, CIRPIN, and TM-vec; smaller ranges (indicating greater invariance to circular permutation) are highlighted in yellow. Plotting the distributions across all synCP combinations reveals each model’s sensitivity to positional reordering. We use the distribution range as a measure of invariance to circular permutation, with smaller ranges reflecting greater invariance, and also report the minimum score to capture cases where a model spuriously indicates dissimilarity due to reordering.

**Supplementary Figure S2:**
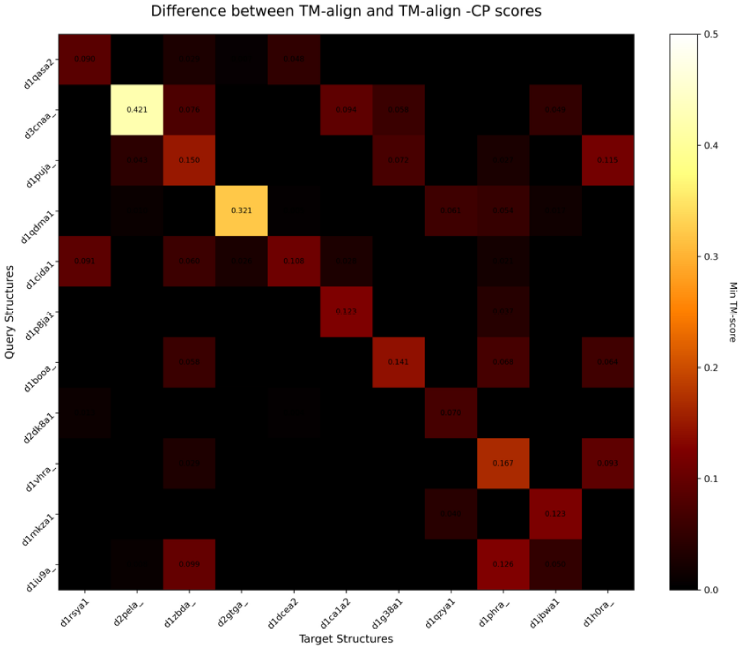
Δ*CP* scores (difference between TM-align and TM-align -cp scores) for the test set of CPs.

**Supplementary Figure S3:**
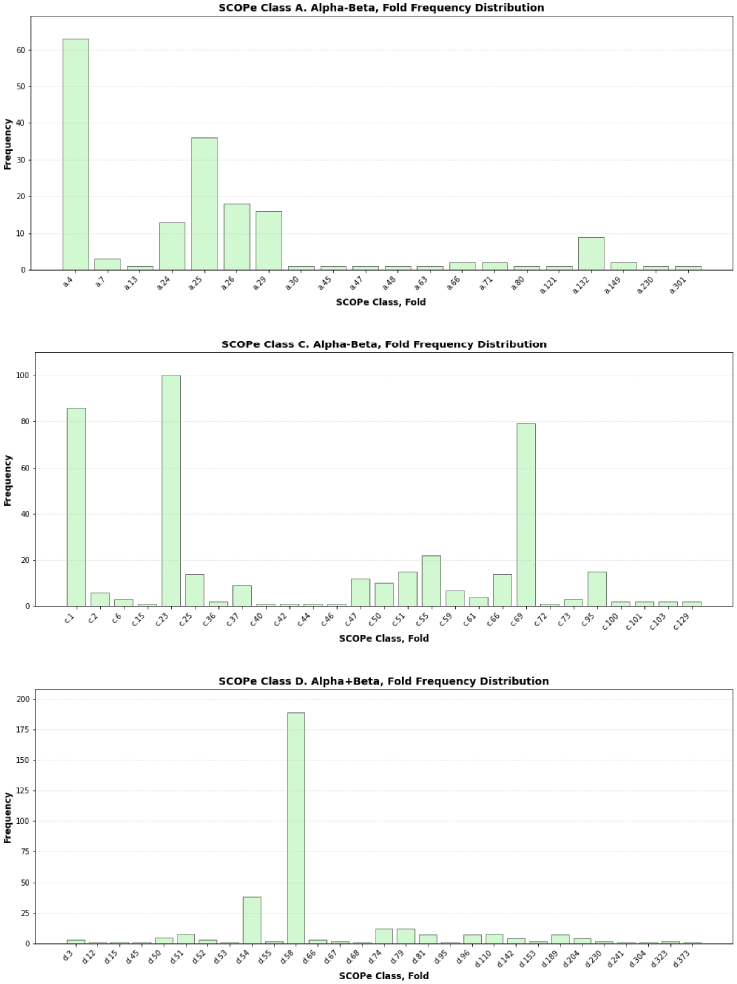
Distribution of SCOPe CPs by SCOPe class and fold

